# Comparative genomics reveals extensive genomic conservation and limited microdiversification among *Xenorhabdus bovienii* isolates recovered from a single *Steinernema feltiae* isolation event

**DOI:** 10.64898/2026.06.04.727993

**Authors:** Cecilia Peralta, Luisa Meier, Leopoldo Palma

**Author notes:** Corresponding author: Leopoldo Palma.

## Abstract

*Xenorhabdus* bovienii is a symbiotic bacterium associated with entomopathogenic nematodes of the genus *Steinernema*. Comparative genomic analyses of closely related isolates provide an opportunity to investigate fine-scale diversification, genome plasticity, and the evolutionary processes shaping symbiotic bacterial populations. Here, we analyzed four *X. bovienii* isolates (XenUTI4.1–XenUTI4.4) recovered from a single *Steinernema feltiae* isolation event using comparative genomics approaches integrating average nucleotide identity (ANI), single-nucleotide polymorphism (SNP) analyses, pangenome reconstruction, biosynthetic gene cluster (BGC) prediction, and mobile element-associated annotation screening.

Whole-genome comparisons revealed extremely high genomic similarity among isolates, with ANI values exceeding 99.84%. Read-based SNP analyses identified only 23–36 annotated variants relative to the XenUTI4.1 reference genome, indicating limited sequence divergence despite detectable microvariation. Functional annotation of these variants showed that most corresponded to missense or synonymous substitutions affecting a small number of coding sequences.

Pangenome analysis identified 4,712 orthologous gene clusters, including a highly conserved core genome of 4,256 clusters (90.3%) shared by all isolates and a relatively small accessory genome comprising 456 clusters. antiSMASH analyses revealed broadly conserved secondary metabolite biosynthetic potential across the four genomes, whereas screening of genome annotations identified abundant phage-related, transposase-associated, and recombination-associated genes consistent with ongoing genome plasticity.

Collectively, these results demonstrate that the analyzed *X. bovienii* isolates represent a highly conserved population exhibiting limited but detectable genomic microdiversification. The coexistence of a large core genome, a modest accessory gene complement, and numerous mobile element-associated functions suggests that localized sequence variation and mobile genetic elements contribute to genomic diversification within *S. feltiae*-associated *X. bovienii* populations.

## 1. Introduction

Entomopathogenic nematodes of the genus *Steinernema* (Rhabditida: Steinernematidae) maintain mutualistic associations with bacteria of the genus *Xenorhabdus*, forming highly specialized symbiotic systems involved in insect pathogenesis and nutrient cycling. During infection, infective juvenile nematodes release their symbiotic bacteria into the insect hemocoel, where the bacteria contribute to rapid host killing, immune suppression, nutrient bioconversion, and protection against competing microorganisms (Nielsen-LeRoux et al., 2012; Palma et al., 2024; Tobias et al., 2017). This mutualistic interaction has attracted considerable interest due to its ecological significance and potential applications in biological pest control (Labaude and Griffin, 2018; Ruiu, 2015).

Among *Xenorhabdus* species, *Xenorhabdus bovienii* is widely associated with multiple *Steinernema* hosts and exhibits substantial genomic and phenotypic diversity. Previous studies have identified variability in virulence-associated factors, secretion systems, toxin production, and biosynthetic gene clusters, suggesting ongoing diversification linked to host adaptation and ecological specialization (Palma et al., 2024; Tobias et al., 2017). Advances in comparative genomics have further enabled strain-level resolution of genomic variation within symbiotic bacterial populations.

Beyond core-genome variation, comparative genomic analyses have increasingly highlighted the importance of accessory genome dynamics, mobile genetic elements, and biosynthetic gene cluster diversification in shaping ecological adaptation among closely related bacterial populations (Medini et al., 2005; Touchon et al., 2009). In *Xenorhabdus* and *Photorhabdus* symbionts, secondary metabolites produced by nonribosomal peptide synthetase (NRPS) and polyketide synthase (PKS) pathways contribute to insect pathogenicity, interbacterial competition, immune suppression, and symbiotic fitness (Tobias et al., 2017). Consequently, integrated analyses combining SNP variation, pangenome structure, secondary metabolite biosynthetic potential, and mobile genetic elements may provide deeper insight into fine-scale diversification processes within symbiotic bacterial populations. Single nucleotide polymorphism (SNP)-based comparative genomics has become a powerful approach for investigating evolutionary relationships among closely related bacterial strains. However, analyses based on draft short-read assemblies remain susceptible to artifacts introduced by repetitive regions, multicopy elements, assembly fragmentation, and ambiguous read mapping (Chen et al., 2020; Wick et al., 2023). Appropriate filtering and variant validation strategies are therefore essential for distinguishing biologically meaningful variation from technical noise.

Despite the growing availability of *Xenorhabdus* genome sequences, fine-scale comparative analyses of *X. bovienii* colonies recovered from the same *Steinernema feltiae* isolation event remain limited. Investigating genomic heterogeneity among colonies originating from a single isolation process may provide insight into within-host diversity, microevolutionary processes, and population structure within symbiotic bacterial communities (Henn et al., 2010; Shapiro and Polz, 2014).

In this study, we analyzed four independently recovered *X. bovienii colonies* (XenUTI4.1–4.4) associated with *S. feltiae* using an integrated comparative genomics framework. By combining ANI analyses, read-based SNP characterization, pangenome reconstruction, biosynthetic gene cluster prediction, and mobile element-associated annotation screening, we sought to characterize fine-scale genomic variation among closely related isolates and evaluate the relative contributions of sequence divergence, accessory genome content, and genome plasticity to diversification within a single symbiotic bacterial population.

## 2. Materials and methods

### 2.1 Isolation of entomopathogenic nematodes and bacterial colonies

Entomopathogenic nematodes were recovered from soil samples collected in Utiel (Valencia, Spain), using *Galleria mellonella* larvae as bait insects. Dead larvae showing symptoms of entomopathogenic nematode infection were transferred to modified White traps to allow emergence and collection of infective juveniles (Palma et al., 2024).

To isolate the bacterial symbionts, infected *G. mellonella* cadavers were surface disinfected with 70% ethanol and haemolymph was streaked onto NBTA agar plates (Akhurst, 1980). Colonies displaying the characteristic morphology of *Xenorhabdus* were subcultured to obtain axenic isolates. Four independently recovered bacterial colonies obtained during the same isolation procedure were selected and designated XenUTI4.1, XenUTI4.2, XenUTI4.3, and XenUTI4.4. Preliminary molecular characterization was performed by PCR amplification of the 16S rRNA gene, and sequencing as previously described for *Xenorhabdus* strains (Palma et al., 2024).

### 2.2 Molecular identification of the nematode

Genomic DNA from infective juveniles (IJs) of the entomopathogenic nematode isolate was extracted using PrepMan™ Ultra Reagent (ThermoFisher Scientific). Briefly, 2–5 IJs were mechanically disrupted in 30 μL of reagent, incubated at 100 °C for 10 min, and centrifuged at 15000 ×*g* for 2 min. The recovered supernatant containing genomic DNA was used as template for PCR amplification of the ITS and D2–D3 regions of the 28S rRNA gene for molecular identification.

The nematode isolate was molecularly identified by amplification of the ITS region and the D2–D3 expansion domains of the 28S rRNA gene. The ITS region was amplified using primers TW81 and AB28, whereas the D2–D3 region was amplified using primers D2A and D3B (Joyce et al., 1994). PCR reactions were carried out in 25 μL volumes containing NZYTaq II 2× Green Master Mix (NZYTech, Portugal), primers, and genomic DNA template.

Amplifications were performed using an initial denaturation step at 94 °C, followed by 35 amplification cycles including denaturation, primer annealing, and extension steps, with a final extension at 72 °C. PCR products were subsequently sequenced at STAB VIDA (Portugal) and compared against reference sequences in GenBank for taxonomic identification of the nematode isolate as *S. feltiae*.

### 2.3 DNA extraction and Illumina sequencing

Bacterial isolates were cultured in LB broth at 28 °C for 48 h with agitation prior to genomic DNA extraction. Total genomic DNA was purified using the Wizard Genomic DNA Purification Kit (Promega, USA) following the manufacturer’s instructions for Gram-negative bacteria. Whole-genome sequencing was performed using Illumina short-read technology at Novogene (Cambridge, UK).

### 2.4 Genome assembly and comparative genomic analyses

Raw Illumina reads were quality-filtered and assembled *de novo* into draft genomes using SPAdes genome assembler v4.2.0 (Bankevich et al., 2012). Assembly quality statistics, including genome size, GC content, contig number, largest contig, and N50 values, were calculated using QUAST (Gurevich et al., 2013). Genome completeness and contamination estimates were evaluated using CheckM (Parks et al., 2015).

Pairwise Average Nucleotide Identity (ANI) analyses were performed using FastANI (Jain et al., 2018) to assess overall genomic similarity among isolates. Initial pairwise SNP comparisons were generated from draft genome assemblies to identify patterns of genomic divergence. To reduce false-positive variant calls associated with repetitive or poorly assembled regions, SNP analyses excluded short contigs and contigs exhibiting anomalously low or extremely high sequencing coverage.

To further investigate fine-scale genomic variation, Illumina reads from each isolate were mapped against the XenUTI4.1 reference genome using Snippy v4.6.0 (https://github.com/tseemann/snippy). High-confidence variants were retained following quality and coverage filtering, and SNP effects were annotated using the corresponding GenBank reference annotation. Variants were subsequently classified according to their predicted effects on coding sequences and used to identify genes affected by microdiversification among isolates.

Genome annotation was performed using the NCBI Prokaryotic Genome Annotation Pipeline (PGAP) and the RAST server (Aziz et al., 2008). Putative biosynthetic gene clusters potentially involved in secondary metabolite production were predicted using antiSMASH (Blin et al., 2019).

Pangenome analyses were performed using Proteinortho v6.3.6 (Lechner et al., 2011) with DIAMOND-based orthology searches to identify shared core-genome and accessory-genome orthologous gene clusters among the four analyzed isolates. Orthologous cluster distributions and genome-sharing patterns were visualized using UpSet plots generated with the UpSetR package in R (Team, 2024).

To explore the contribution of genome plasticity and horizontal genomic dynamics, RAST-derived genome annotations were screened for mobile genetic element-associated terms, including phage-related proteins, transposases, insertion sequence-associated proteins, integrases, recombinases, and resolvases. Annotation counts were subsequently compared among genomes to evaluate variation in mobile element-associated gene content.

Species assignment and genome-based taxonomic analyses were performed using the Type (Strain) Genome Server (TYGS) (Meier-Kolthoff and Goker, 2019) and PubMLST (Jolley et al., 2018).

Plots and statistical analyses were generated using the R packages ggplot2, ape, pheatmap, dplyr, and ggrepel and UpSetR.

## 3. Results

### 3.1 Nematode identification, bacterial genome assembly statistics and general genomic features

Molecular identification based on ITS and 28S rRNA (D2–D3) sequences consistently assigned the nematode isolate to *S. feltiae*. The ITS sequence showed 99.13% identity to *S. feltiae* strain A2 (AY230170.1), while the 28S rRNA sequence showed 98.55% identity to *S. feltiae* isolate SFQ16 (OR651369.1), confirming species-level identification.

Draft genome assemblies of the four *X. bovienii* isolates revealed generally similar genome characteristics, with only minor differences in assembly continuity and genome size (Table 1). Genome sizes ranged from 4.59 Mb (XenUTI4.3) to 5.41 Mb (XenUTI4.2), whereas GC content varied only slightly, between 44.40% and 47.76%.

**Table 1.** General genome assembly statistics of *X. bovienii* isolates.

| Isolate | Size (bp) | GC (%) | Contigs | Largest contig (bp) | N50 (bp) | L50 | Completeness (%) |
| --- | --- | --- | --- | --- | --- | --- | --- |
| XenUTI4.1 | 4,571,669 | 44.5 | 295 | 79,896 | 31,047 | 50 | 100 |
| XenUTI4.2 | 4,569,791 | 44.5 | 259 | 108,917 | 32,449 | 44 | 100 |
| XenUTI4.3 | 4,592,433 | 44.7 | 447 | 78,641 | 23,074 | 63 | 100 |
| XenUTI4.4 | 4,667,578 | 44.4 | 264 | 145,843 | 43,727 | 36 | 100 |

Assembly statistics were generated using QUAST. Completeness metrics were obtained with CheckM.

Among the analyzed isolates, XenUTI4.3 exhibited a somewhat smaller assembly size and lower N50 value than the remaining genomes. However, overall assembly statistics remained broadly comparable among isolates and did not affect the high levels of genomic similarity observed in subsequent comparative analyses.

QUAST analyses revealed moderate variation in contig distribution among assemblies, with XenUTI4.3 containing fewer large contigs and a lower cumulative assembly size than the remaining genomes. Nevertheless, these assembly-level differences contrasted with the strong genomic conservation revealed by ANI, SNP, and pangenome analyses.

Pairwise Average Nucleotide Identity (ANI) analyses confirmed that all isolates belonged to the same *X. bovienii* genomic lineage, displaying ANI values above 99.91% across all comparisons (Table 2). The highest ANI values were observed between XenUTI4.1 and XenUTI4.2 (99.97%) and between XenUTI4.3 and XenUTI4.4 (99.97%), indicating extremely limited genomic divergence among isolates.

**Table 2.** Pairwise Average Nucleotide Identity (ANI) values among *X. bovienii* isolates.

|  | XenUTI4.1 | XenUTI4.2 | XenUTI4.3 | XenUTI4.4 |
| --- | --- | --- | --- | --- |
| XenUTI4.1 | 100 | 99.97 | 99.94 | 99.92 |
| XenUTI4.2 | 99.97 | 100 | 99.93 | 99.91 |
| XenUTI4.3 | 99.94 | 99.93 | 100 | 99.97 |
| XenUTI4.4 | 99.92 | 99.91 | 99.97 | 100 |

ANI values above 99.87% confirmed that all isolates belong to the same *X. bovienii* genomic lineage. XenUTI4.1 and XenUTI4.2 formed a closely related pair (99.97% ANI), while XenUTI4.3 and XenUTI4.4 formed a second closely related pair (99.97% ANI).

### 3.2 Pairwise SNP analyses and filtering of low-confidence regions

Read-based SNP analysis using the cleaned XenUTI4.1 assembly as reference revealed extremely limited nucleotide variation among the four *X. bovienii* isolates. Variant counts relative to XenUTI4.1 ranged from 18 to 35 high-confidence SNPs, with XenUTI4.3 displaying the fewest variants and XenUTI4.2 the highest. Pairwise SNP distances ranged from 18 to 35 variants, indicating only minor sequence divergence among isolates. Overall, these results demonstrate a high degree of nucleotide-level conservation among isolates independently recovered during the same *S. feltiae* isolation event (Figure 1).

**Figure 1.**
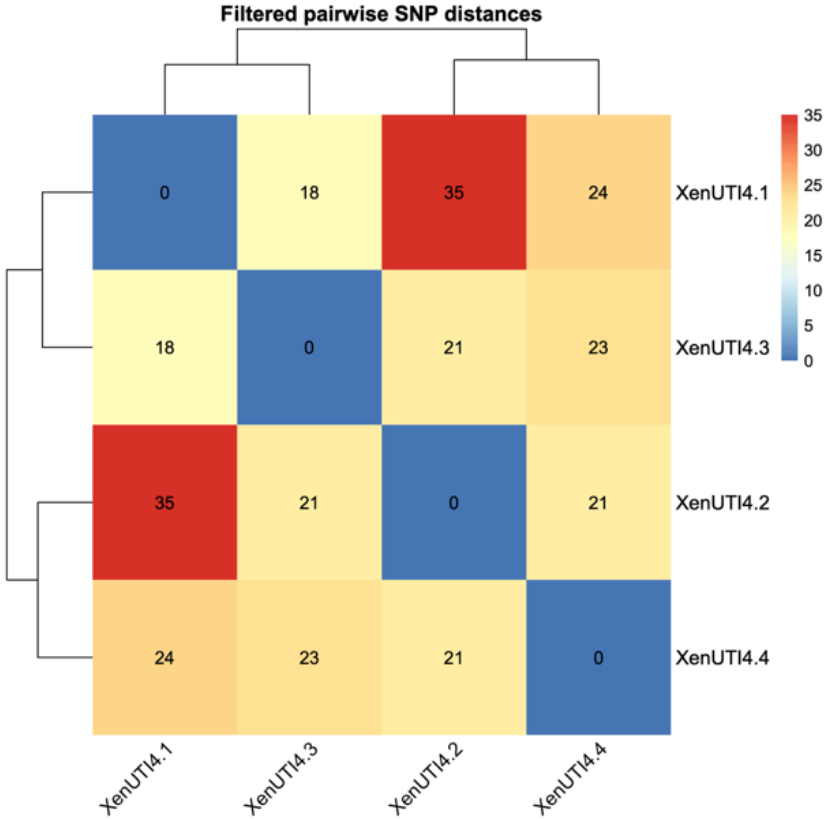
Pairwise SNP distance heatmap among *X. bovienii* isolates. Distances were calculated from high-confidence SNPs identified by read-based mapping against the XenUTI4.1 reference genome. All pairwise comparisons revealed low SNP counts (18–35 variants), consistent with the high genomic similarity observed among isolates recovered from the same *S. feltiae* isolation event.

### 3.3 Multivariate analyses

Principal component analysis (PCA) based on pairwise SNP distances revealed limited genomic differentiation among the four *X. bovienii* isolates (Figure 2). The ordination was consistent with the low SNP counts observed across all pairwise comparisons and did not identify any strongly differentiated lineage. Together with the ANI and SNP-distance analyses, these results indicate the presence of limited genomic microvariation within an otherwise highly conserved *X. bovienii* population.

**Figure 2.**
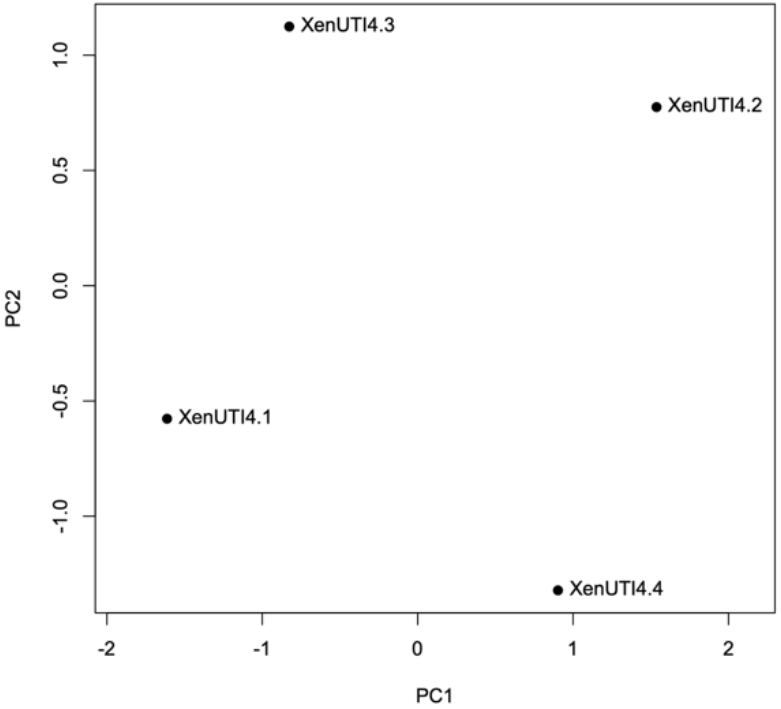
Principal component analysis (PCA) of pairwise SNP distances among *X. bovienii* isolates. PCA was performed using the matrix of pairwise SNP distances obtained from read-based variant calling. The ordination reflects the limited genomic variation detected among isolates and is consistent with their overall high genomic similarity.

### 3.4 Virulence-associated genes and toxin-related proteins

BLASTX screening against the BPPRC database identified a conserved set of toxin-associated homologs across all four *X. bovienii* isolates, including App1Ba1-, Mcf1Ab1-, and Mcf1Ba1-related proteins (one homolog each), as well as additional *Photorhabdus*-associated proteins. No qualitative differences in the presence or absence of these homologs were detected among isolates.

Genome annotation using RAST similarly revealed highly conserved repertoires of virulence-associated genes, including putative insecticidal toxin complex proteins (7 copies), probable insecticidal toxins (12 copies), chitin-binding proteins (1 copy), extracellular serine protease precursors (2 copies), RTX toxin transporters (2 copies), and regulatory components such as PhoQ (1 copy) and VirK (1 copy). Type IV secretion system-associated proteins were also consistently identified across all genomes (3 copies).

Overall, virulence- and toxin-associated gene repertoires were highly conserved among isolates, supporting the extensive genomic conservation revealed by ANI, SNP, and pangenome analyses.

### 3.5 Functional genome annotation and metabolic categories

Comparative functional annotation based on RAST revealed highly similar coding potential among the four *X. bovienii* isolates (Figure 3). All genomes displayed comparable representation of genes associated with metabolism, transport systems, regulation and signaling, secretion and export pathways, stress response mechanisms, motility-related functions, cell wall and membrane-associated processes, and virulence-associated categories.

**Figure 3.**
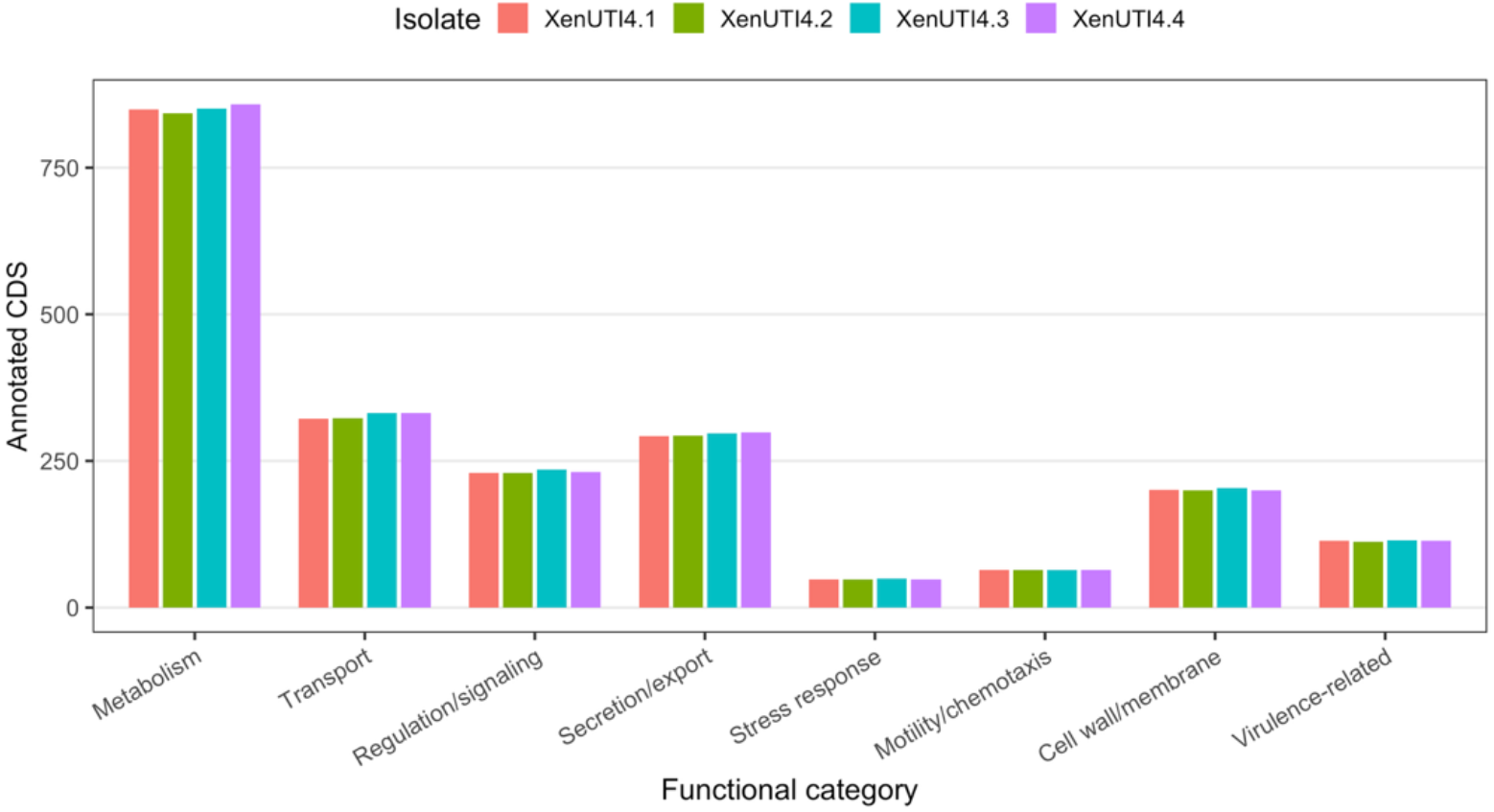
Comparative distribution of RAST-derived functional annotation categories among *X. bovienii* isolates. Bar plots represent the number of annotated coding sequences assigned to major functional categories. Similar profiles across all genomes indicate a highly conserved metabolic and physiological repertoire.

Metabolism-associated functions represented the largest category in all genomes, followed by transport-, regulation-, and secretion-related functions. Differences among isolates were minimal across all functional categories, with only minor variation in gene counts and no evidence of isolate-specific functional enrichments. Motility- and chemotaxis-related functions showed nearly identical representation across all genomes.

Overall, the functional profiles indicate a highly conserved metabolic and physiological repertoire among the analyzed *X. bovienii* isolates, consistent with the high ANI values, limited SNP divergence, and extensive core-genome conservation revealed by comparative genomic analyses.

### 3.6 Fine-scale SNP annotation and intra-isolation divergence

To further characterize the limited genomic microdiversification detected among isolates, cleaned Illumina reads were mapped against the XenUTI4.1 reference genome and variants were functionally annotated. A total of 36, 23, and 24 variants were identified in XenUTI4.2, XenUTI4.3, and XenUTI4.4, respectively, relative to XenUTI4.1.

Most variants occurred within coding sequences and included both synonymous and missense substitutions. Recurrent missense variants were detected in genes associated with iron acquisition, secondary metabolism, mobile genetic elements, and phage-related functions. Affected loci included the ferric enterobactin transport permease FenD, polyketide synthase-associated proteins, phage tail fiber proteins, conjugative transfer proteins, and mobile element-associated proteins, among others. In addition, a missense variant was identified in a VirG protein associated with type VI secretion systems.

Functional classification of genome variants identified relative to the XenUTI4.1 reference genome revealed that most substitutions were located within coding regions and were predominantly missense or synonymous mutations (Table 3).

**Table 3.** Functional classification of genomic variants identified relative to the XenUTI4.1 reference genome. Variants were categorized as missense, synonymous, complex/MNP, or non-coding according to their predicted functional effects. Variant counts are shown for XenUTI4.2, XenUTI4.3, and XenUTI4.4 relative to the XenUTI4.1 reference genome.

| Isolate | Total variants | Missense | Synonymous | Complex/MNP | Non-coding |
| --- | --- | --- | --- | --- | --- |
| XenUTI4.2 | 36 | 16 | 11 | 8 | 1 |
| XenUTI4.3 | 23 | 10 | 6 | 5 | 2 |
| XenUTI4.4 | 24 | 10 | 7 | 6 | 1 |

XenUTI4.2 exhibited the highest number of annotated variants, whereas XenUTI4.3 and XenUTI4.4 showed lower levels of divergence. Collectively, these results indicate that genomic variation among isolates is limited and concentrated in a small number of coding loci, consistent with the extensive genomic conservation observed throughout the study.

### 3.7 Biosynthetic gene cluster diversity

Comparative analyses revealed highly conserved biosynthetic gene cluster (BGC) repertoires across all four *X. bovie*nii isolates. Detected clusters were dominated by non-ribosomal peptide synthetases (NRPSs), accompanied by NRPS-like systems, type I polyketide synthases (T1PKS), PKS-associated loci, polyunsaturated fatty acid (PUFA) biosynthetic regions, siderophore-associated clusters, and additional secondary metabolite biosynthetic systems.

Although minor differences in cluster composition were observed, overall BGC repertoires remained highly similar among isolates. XenUTI4.3 contained the largest number of predicted BGCs (35), including one additional T1PKS-associated cluster, whereas XenUTI4.2 contained a trans-AT-PKS-associated region not detected in the remaining genomes. In contrast, XenUTI4.4 displayed a slightly reduced BGC repertoire (29 clusters) relative to the other isolates.

Nevertheless, all strains retained a largely shared complement of secondary metabolite biosynthetic loci, indicating extensive conservation of biosynthetic potential despite the limited genomic divergence observed among isolates (Figure 4, Table 4).

**Table 4.**
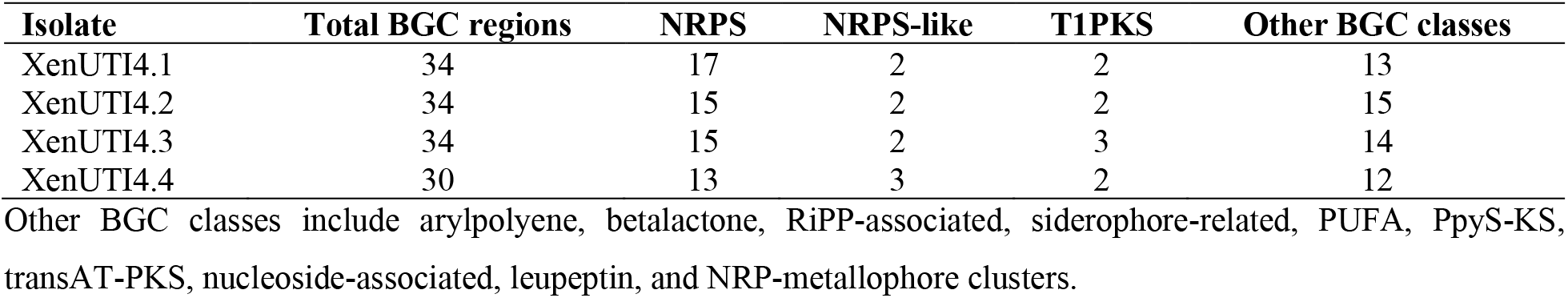
Summary of antiSMASH-predicted biosynthetic gene clusters in *X. bovienii* isolates.

| Isolate | Total BGC regions | NRPS | NRPS-like | T1PKS | Other BGC classes |
| --- | --- | --- | --- | --- | --- |
| XenUTI4.1 | 34 | 17 | 2 | 2 | 13 |
| XenUTI4.2 | 34 | 15 | 2 | 2 | 15 |
| XenUTI4.3 | 34 | 15 | 2 | 3 | 14 |
| XenUTI4.4 | 30 | 13 | 3 | 2 | 12 |

Other BGC classes include arylpolyene, betalactone, RiPP-associated, siderophore-related, PUFA, PpyS-KS, transAT-PKS, nucleoside-associated, leupeptin, and NRP-metallophore clusters.

**Figure 4.**
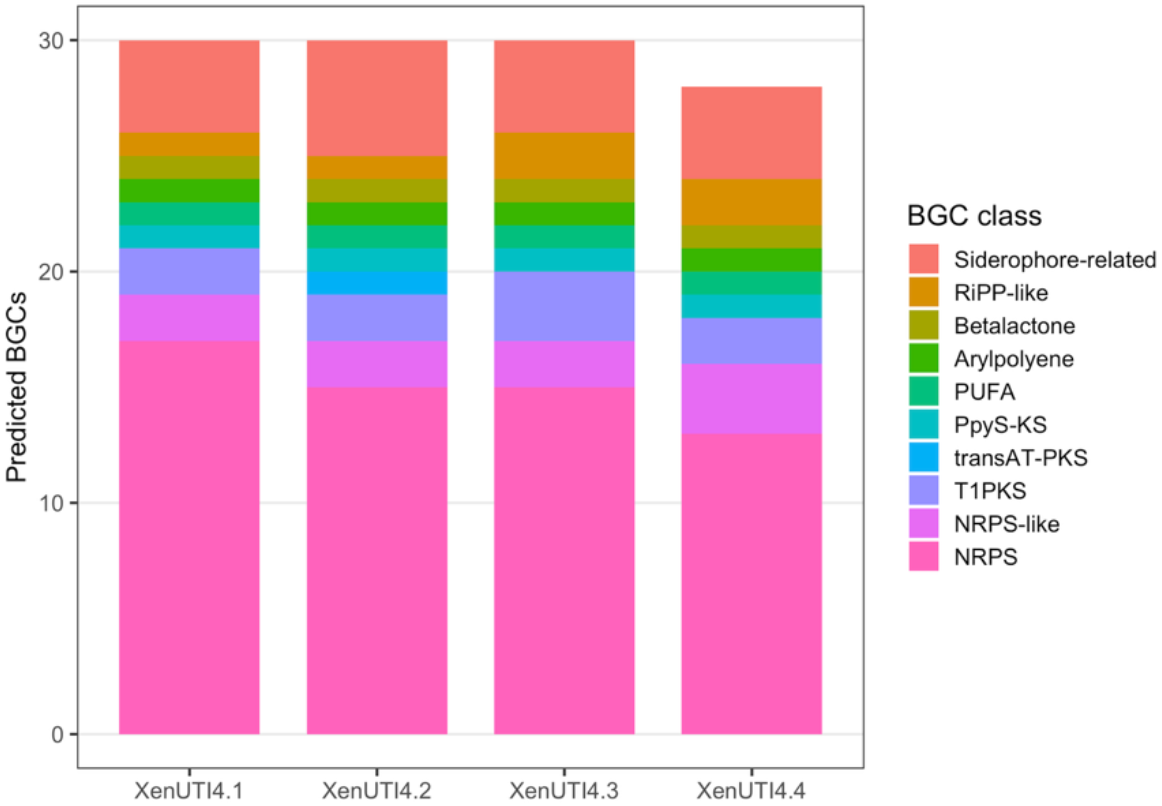
Comparative distribution of antiSMASH-predicted biosynthetic gene clusters (BGCs) among *X. bovienii* isolates. Stacked bar plots summarize the abundance of major BGC classes identified in each genome. Non-ribosomal peptide synthetase (NRPS) clusters represented the predominant biosynthetic category across all isolates, followed by NRPS-like, polyketide synthase (PKS)-associated, siderophore-related, and RiPP-associated clusters. Overall BGC repertoires were highly conserved across isolates. Minor differences in cluster composition were observed, with XenUTI4.3 displaying the largest predicted repertoire and XenUTI4.4 showing a slightly reduced number of clusters.

### 3.8 Pangenome structure and accessory genome composition

Comparative pangenome analysis of the four *X. bovienii* isolates identified 4,712 orthologous gene clusters. Of these, 4,256 clusters (90.3%) were shared by all isolates and constituted the core genome, whereas 456 clusters formed the accessory genome. A total of 4,417 clusters (93.7%) were present in at least three isolates, further indicating extensive conservation of gene content across the collection (Figure 5).

**Figure 5.**
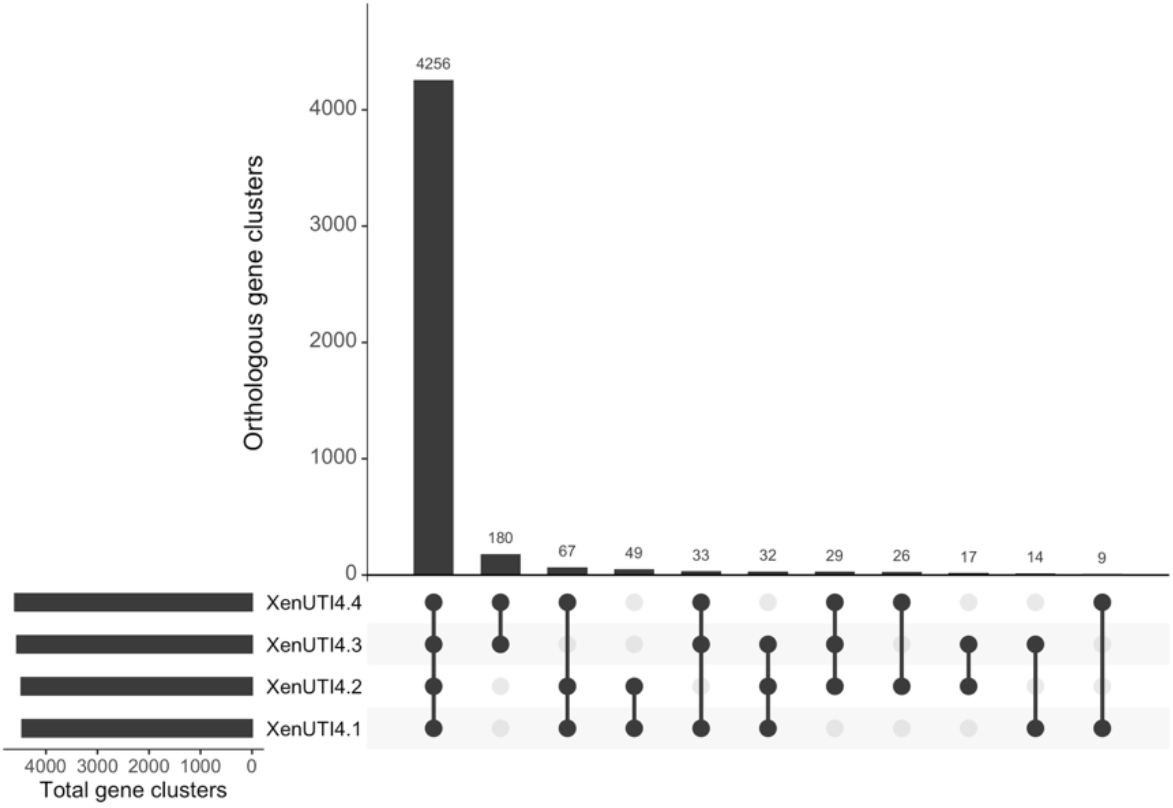
UpSet plot of orthologous gene cluster distribution among the four *X. bovienii* isolates. Proteinortho analysis identified 4,712 orthologous gene clusters, of which 4,256 were shared by all isolates and constituted the core genome. The remaining 456 clusters formed the accessory genome and were distributed among individual isolates or subsets of genomes.

Pairwise orthology comparisons further supported the high level of genomic similarity among isolates. The greatest number of shared orthologous clusters was observed between XenUTI4.1 and XenUTI4.2, consistent with their high ANI values and overall genomic similarity. Although XenUTI4.3 and XenUTI4.4 displayed slightly fewer shared clusters with the remaining isolates, overall gene-content differences were limited and involved only a small fraction of the total pangenome.

Overall, pangenome analysis demonstrated that the four isolates possess a highly conserved genomic backbone while retaining a relatively small accessory genome that contributes to fine-scale strain-level variation. These findings are consistent with the high ANI values, limited SNP divergence, conserved functional profiles (and broadly similar biosynthetic gene cluster repertoires observed across the analyzed isolates (Figure 5).

### 3.9 Mobile element-associated annotations

Screening of RAST-derived genome annotations for mobile genetic element-associated terms revealed abundant phage-related, transposase/mobile element, and recombinase/integrase/resolvase annotations across all analyzed genomes. Phage-associated annotations ranged from 194 to 220 genes per genome, whereas transposase and mobile element-associated annotations represented the most abundant category, ranging from 861 to 987 genes. Recombination-associated annotations, including integrases, recombinases, and resolvases, were also detected in all isolates, with counts ranging from 40 to 46 genes.

XenUTI4.3 and XenUTI4.4 exhibited slightly higher numbers of phage-associated, mobile element-associated, and recombination-related annotations than XenUTI4.1 and XenUTI4.2, although all genomes contained substantial repertoires of these functions.

Although these analyses do not constitute formal prophage or genomic island prediction, the widespread occurrence of mobile element-associated annotations suggests that horizontal gene transfer and genome plasticity have contributed to genome evolution within the analyzed *X. bovienii* population. These observations are consistent with the accessory genome identified by pangenome analysis and may partially explain the limited gene-content variation observed among isolates despite their highly conserved core genome (4,256 of 4,712 orthologous gene clusters). The abundance of transposases, recombinases, and phage-associated functions further suggests ongoing genomic remodeling processes that could facilitate the acquisition, loss, or rearrangement of accessory genetic material and contribute to the fine-scale microdiversification observed among isolates.

## 4. Discussion

Comparative genomic analyses of the four *X. bovienii* isolates revealed detectable genomic heterogeneity despite their recovery from the same *S. feltiae* isolation event. Although all genomes displayed extremely high ANI values (>99.84%), read-based SNP analyses identified a limited number of high-confidence variants among isolates, indicating the presence of fine-scale genomic microdiversification within an otherwise highly conserved population. These findings illustrate how closely related symbiotic bacteria can retain detectable evolutionary variation despite exhibiting near-identical genome-wide nucleotide identity (Henn et al., 2010; Shapiro and Polz, 2014).

An important observation of this study was the substantial reduction in apparent genomic variation following the removal of low-confidence regions during SNP analyses. Initial variant detection identified considerably higher levels of polymorphism; however, many of these variants were associated with short contigs, repetitive regions, or genomic regions exhibiting anomalous sequencing coverage. Filtering these regions markedly reduced SNP counts while preserving overall relationships among isolates. These results emphasize the importance of stringent quality-control procedures when interpreting genomic variation from short-read draft assemblies, particularly in bacterial genomes containing repetitive sequences, multicopy elements, and mobile genetic elements (Chen et al., 2020; Wick et al., 2023).

Although minor differences in genome size, assembly continuity, and SNP content were observed among isolates, all comparative analyses consistently supported a high degree of genomic conservation. The low number of annotated SNPs detected relative to the reference genome, together with ANI values exceeding 99.84%, indicates that the four isolates likely represent closely related members of the same bacterial population. Rather than supporting the existence of strongly differentiated lineages, the results are more consistent with ongoing microevolutionary processes generating localized genomic variation within a largely conserved symbiotic population.

The conservation observed at the nucleotide level was mirrored by the overall preservation of functional gene repertoires. Comparative annotation analyses indicated that genes associated with metabolism, transport, regulation, stress response, and secretion systems were broadly conserved across all genomes. Likewise, virulence-associated functions commonly reported in *Xenorhabdus* species, including toxin-related proteins, extracellular enzymes, secretion-associated factors, and other insect-pathogenicity-associated determinants, were consistently represented in all isolates. These observations suggest that the core biological functions required for symbiosis, insect pathogenicity, and environmental persistence remain highly conserved despite the presence of detectable genomic microvariation (Nielsen-LeRoux et al., 2012; Tobias et al., 2017).

Pangenome reconstruction further reinforced the remarkable genomic similarity among isolates. A total of 4,712 orthologous gene clusters were identified, of which 4,256 (90.3%) constituted a conserved core genome shared by all four isolates. Only 456 clusters formed the accessory genome, indicating that gene content variation among isolates was relatively limited. The predominance of a large, shared core genome is consistent with the high ANI values and low SNP counts observed throughout the study and suggests that diversification within this population is driven primarily by localized sequence variation and a comparatively small accessory gene complement rather than extensive gene gain or loss. Similar patterns have been reported in closely related bacterial populations in which microdiversification occurs against a background of strong genomic conservation (Medini et al., 2005; Shapiro and Polz, 2014).

Analysis of biosynthetic gene clusters revealed broadly conserved secondary metabolite biosynthetic potential across the four genomes. Most predicted NRPS-, PKS-, and hybrid biosynthetic gene clusters were shared among isolates, indicating that the principal secondary metabolite repertoire characteristic of *X. bovienii* remains stable within this population. In *Xenorhabdus* and *Photorhabdus* symbionts, secondary metabolites play important roles in insect virulence, interbacterial competition, immune suppression, and maintenance of the nematode-bacterium association (Tobias et al., 2017). The overall conservation of biosynthetic gene cluster repertoires observed here suggests that these ecological functions are maintained across isolates despite limited strain-level genomic variation.

In contrast, mobile element-associated annotations were abundant in all genomes. Numerous phage-related proteins, transposases, integrases, recombinases, and resolvases were identified, indicating that horizontal gene transfer and genome plasticity likely contribute to the evolutionary dynamics of these bacteria. The slightly higher abundance of mobile element-associated annotations detected in XenUTI4.3 and XenUTI4.4 compared with XenUTI4.1 and XenUTI4.2 may partially explain the modest differences observed in accessory gene content. Together, these findings suggest that mobile genetic elements may play a central role in generating and maintaining genomic variability within otherwise highly conserved *X. bovienii* populations.

Such discrepancies likely reflect the fact that SNP analyses primarily capture variation within conserved genomic regions, whereas k-mer-based approaches may also incorporate signals derived from accessory genome composition and repetitive sequences. Similar differences between core-genome and accessory-genome evolutionary signals have been reported in other bacterial comparative genomic studies and highlight the complexity of diversification processes operating within closely related microbial populations (Wick et al., 2023).

Several limitations should be acknowledged. All analyses were performed using Illumina short-read draft assemblies, which remain susceptible to assembly fragmentation and incomplete reconstruction of repetitive genomic regions, mobile elements, and complex biosynthetic loci. Consequently, some aspects of genome architecture and accessory genome composition may remain unresolved. The incorporation of long-read sequencing technologies and hybrid assembly approaches will enable more accurate reconstruction of chromosome structure, mobile genetic elements, and biosynthetic gene clusters, thereby providing improved resolution of evolutionary relationships within this population.

To our knowledge, this study represents one of the first integrated comparative genomic analyses examining multiple *X. bovienii* colonies recovered from a single *S. feltiae* isolation event. The coexistence of a highly conserved core genome, limited accessory genome variability, and abundant mobile element-associated functions highlights the dynamic balance between genomic stability and diversification within symbiotic bacterial populations. These findings provide a foundation for future investigations into the microevolutionary processes, ecological adaptation, and genome plasticity that shape the evolution of Xenorhabdus symbionts.

## 5. Conclusion

This study provides a fine-scale comparative genomic analysis of four *X. bovienii* isolates independently recovered from a single *S. feltiae* isolation event. Despite extremely high overall genomic similarity, complementary analyses based on ANI, SNP characterization, pangenome reconstruction, biosynthetic gene cluster prediction, and mobile element-associated annotations revealed detectable genomic microdiversification among isolates.

The analyzed genomes shared a highly conserved core genome comprising 4,256 orthologous gene clusters (90.3% of the pangenome), while only a limited accessory gene complement contributed to strain-level variation. Biosynthetic gene cluster repertoires were largely conserved across isolates, whereas abundant mobile element-associated functions suggested ongoing genome plasticity and potential horizontal gene transfer.

Collectively, these findings indicate that diversification within this *X. bovienii* population is characterized by localized sequence variation and modest accessory genome dynamics rather than extensive genomic divergence. This work provides a genomic framework for future studies investigating microevolutionary processes, genome plasticity, and ecological adaptation in *Xenorhabdus* symbionts.

## CRediT authorship contribution statement

Conceptualization, L.P., methodology, C.P., L.M. and L.P.; formal analysis, L.P.; investigation, C.P., L.M. and L.P.; resources, L.P.; data curation, L.P.; writing—original draft preparation, L.P.; writing, review and editing, C.P. and L.P.; visualization, L.P.; supervision, L.P.; project administration, L.P.; funding acquisition, L.P. All authors have read and agreed to the published version of the manuscript.

## Declarations

### Ethical approval

This article does not contain any studies involving humans performed by any of the authors. Experiments involving nematodes were conducted in accordance with institutional and national guidelines applicable to invertebrate model organisms.

### Availability of data

Genome assemblies generated in this study have been submitted to the NCBI GenBank database under BioProject PRJNA1439957. Individual accession numbers for the four *X. bovienii* isolates will be incorporated upon completion of the submission and annotation process.

## Acknowledgements

Leopoldo Palma gratefully acknowledges the Spanish Ministry of Science, Innovation, and Universities, the Spanish State Research Agency, and the European Union for funding his Ramón y Cajal contract (grant ref. RYC2023-043507-I).

